# Long-range infra-sound acoustic signaling in human *in vivo*

**DOI:** 10.1101/233353

**Authors:** Xiaoyue Chang, Li Zhang, Xiguang Wei, Zengfu Peng, Zhiguo Zhang, Zhong Liu, Pei-Wen Li, Pui Tao Lai, Shing Chow Chan, Edward S Yang

**Affiliations:** University of Hong Kong. Hong Kong SAR, China; Hong Kong Baptist University. Hong Kong SAR, China; Shenzhen University. China; National Chiao-Tung University. Taiwan; Columbia University. United States of America

## Abstract

Acupuncture is widely deployed today, but its basic physiology with Qi and meridian is not understood. This letter postulates that Qi is an infrasound wave packet and meridian is the muscle waveguide. Using video cameras and signal processing in an IRB approved clinical experiment we performed a comparison between control and electro-activated statistical tests on the long range (50-80cm) unidirectional transmission of Qi (p = 0.025). In the reverse direction, there is no transmission even in a distance of less than 10 cm (p = 0.545). The rectification is a surprise but in full agreement with Huang Di Nei Jing.

---

This article presents clinical evidence of Qi and Meridian, two mystical unknowns that are described extensively in the ancient Chinese Medicine literature and accepted by all Chinese medicine practitioners but have never been observed or measured. The best description of Qi and meridian is given in Huang Di Nei Jing (2nd Century BC) of which Unschuld has given us two superb translations in Su Wen & Ling Shu (1, 2). Many scientists, both Chinese and non-Chinese, question the scientific principles given in these ancient medical texts (3).But to the translator, these historical classics were, quoted here, “revolutionary work in the true sense of the word. They confronted their contemporaries with the idea that natural laws were valid regardless of deities, spirits, demons and ancestors, as well as time and space. These intellectuals constituted the kernel of an enlightened, secular perspective on the world, which would open up a view of nature and the embedding of humanity in the laws of nature as a foundation for understanding the origins, essence and transience of life. The concept of natural phenomena obeying laws was new and certainly disturbing to the majority of people for whom the existence of deities, spirits, demons and ancestors seemed perfectly obvious. The groundwork was thereby laid for a completely new form of healing arts we call ‘medicine’. The intellectuals had removed health and sickness from the religious view of the world. They regarded the laws of nature as the sole standard that decided the well- or ill-being of each individual…They cited statistics that reflect a time span of experience with many patients…An important new concept stood for these resources: qi, the pictogram formed in the later Zhou or early Han period when the new medicine was also being conceived, refers to vapors of the most minute materials that, along with blood, are essential for the survival of the human body”[Introduction in Ling Shu (2)]

“The twelve conduits (meridians) transmit much or little blood and little or much qi, all these can be quantified.” [Chapter 2, Ling Shu (2)]

“In both ancient Greek and Chinese medicine, the concept of “blood” as a visibly red bodily fluid contrasted with that of the invisible yet equally essential vapors. Greeks called these pneuma and referred to the arteries literally translated as “air vessels”, since the arteries appear empty when a corpse is dissected. The authors of the ancient Chinese texts regarded the matter somewhat differently. Only the blood is bound to vessels, they thought. On the other hand, vapors or qi can make their way anywhere in the body. They enter the body through food and drink, as well as via the air inhaled through the mouth and nose. They can leave the body through the mouth and nose, through the skin and as flatulence.” [Introduction in Ling Shu (2)]

There is a great deal of misunderstanding of Chinese medicine in the general public as well as the science community. These Nei Jing quotes are added to show the precise understanding of natural laws 2000 years ago, a revelation to those of us who come across it the first time. But most of the modern scientists are so absolutely sure that there is a religious fervor in rendering a judgment. Although the ancient physicians did not know what science is, they had placed clinical evidence as their only measuring yardstick even though they did not know how to measure Qi and meridians. Our desire of measuring them is to clarify the laws of nature our ancestors found as clinical phenomena but lacking the science to expound and elucidate it quantitatively. The missing science apparently has not stopped its implementation as an art of healing.

To the non-specialists, the Nei Jing deals with clinical evidence within the context of ancient abstract Chinese worldviews, making it difficult to separate medicine from philosophy (3). Acupuncture is found effective (4–8) and broadly deployed today but its physiology is a puzzle not understood. The missing element appears to be science. While it has millions of users, there are not a small number of non-believing scientists who denigrate it and demand scientific verification. To validate these classics in our study, we postulate Qi and Meridian as measurable substance and accept the Nei Jing as a text of clinical medicine but excluding its philosophy. To put it simply, our objective is to bring science back into the art of needling.

### Significance Statement

It appears we have discovered Qi (pneuma) as a measurable human body signal using Meridians (muscle tissue conduits) as a unidirectional transmission network. The experiment was performed in 12 human volunteers’ *in vivo* using acupuncture as the basic technique. Low frequency electrical pulses were applied to two needles converting them to mechanical vibrations in the back of the shoulder as drivers and 3 needles are inserted at acupoints along the meridians in the back of the neck, at the right elbow and in the back of the right hand respectively. The far away needles (50 and 80 cm) have strong resonance down streams but not the nearby needle (10 cm) in the reverse direction. Data confirm Huang Di Nei Jing.

Let us follow the physical description given in Nei Jing. The first thing we found is, without ambiguity, that the signaling in acupuncture does not make use of nerves. In fact, it is through the use of a material parameter called Qi, passing through a network called Jing-luo (Meridians). We are told that Qi is a finest matter existing in all possible aggregate states, from air, steam, vapor, liquid, and even solid matter. Nei Jing further states that for a normal person, Qi is generated from inhale and exhale and it changes internally to move and emerge (1). Using current vernacular, Qi appears to reside in matter of the gas phase, liquid phase or solid state and delivers a twoway motion. Not visible or audible, Qi is a time dependent phenomenon produced by cyclical mechanical function and capable to transform internally, e.g. from breathing motion and its frequency to that of heart-beat and blood flow. It can be sensed indirectly by change in outward appearance or felt by touch. Taking breathing as a start, Nei Jing seems to have described an infra-sound wave packet, not a mystical spirit or indescribable energy. In other words, the physical properties of Qi appear to be in complete agreement with science if one stays in medicine and away from philosophical speculations. Meridian then is the muscle waveguide for the transmission of the wave packet Qi. Mechanical movement was well known in the time of Nei Jing (5 centuries or more after Psalm 119:120’s depiction of trembling muscle). Qi was recognized as a moving signal initiated by breathing and propagate in all parts of the body through gas, liquid or solid materials. In Nei Jing's description, Qi appears not just a mechanical or acoustic wave packet; it must have incorporated molecules including oxygen, water, carbon dioxide, adenosine and calcium ions making it a fundamental part of a living being (9).

To test the foregoing hypothesis, an experimental study was carried out in the HKBU Acupuncture Clinic with approved IRB (HASC/13-14/0172) by Human and Animal Studies Committee of Hong Kong Baptist University and participant consent signed. Healthy volunteers, 4 men and 8 women, aged between 22-32 with no history of acupuncture treatment, chronic diseases, medication or pains were recruited. An electro-acupuncture Unit (ES-160, IT0 Co. Ltd, Tokyo, Japan) was used to deliver pulses with frequency of 3.3 Hz and 10Hz, alternating in a 3-sec cycle on two driver needles. Other needles without electricity, are properly placed on a recognized meridian to serve as distant sensors. The needle vibrations are captured by high-speed cameras before converting to frequency spectra via Fourier transform. (See Method Section) The volunteers, acupuncturists and data processing scientists do not know and cannot predict the results so the clinical experiment is double-blind.

In order to study the mechanical wave packet signaling and its transmission, volunteers were tested to compare theobserved needle vibrations with and without electrical stimulation. The Large Intestine (LI) Meridian of Hand-Yang Ming (10, 11) was selected as the main conduit with needle placements from the back-shoulder-neck to the elbow and hand. A layout of the needles on the right hand is shown in Figure 1 where needles #2-#3 are used as driver (with electricity) connections: #2-GB21 and #3-GV14, both are acupoints on meridians other than LI. In addition, three sensing needles are inserted: #1 at acupoint BL10 on the back side of the neck about 10 cm, #4 at acupoint LI11 at the elbow about 50 cm and #5 at acupoint LI4 on the back side of the right hand about 80 cm away from #2 and #3 (see Figure 1). The design has implications that one collateral and four meridians, the large-intestine, the bladder, the gallbladder and the governor vessel (all originated in Nei Jing) are being tested not as isolated conduits but as an interconnecting network. The needle penetration depth is between 1.5 and 2 cm. The needle placement and depth are clinically typical and no attempt was made to be precisely duplicated.

**Fig. 1.**
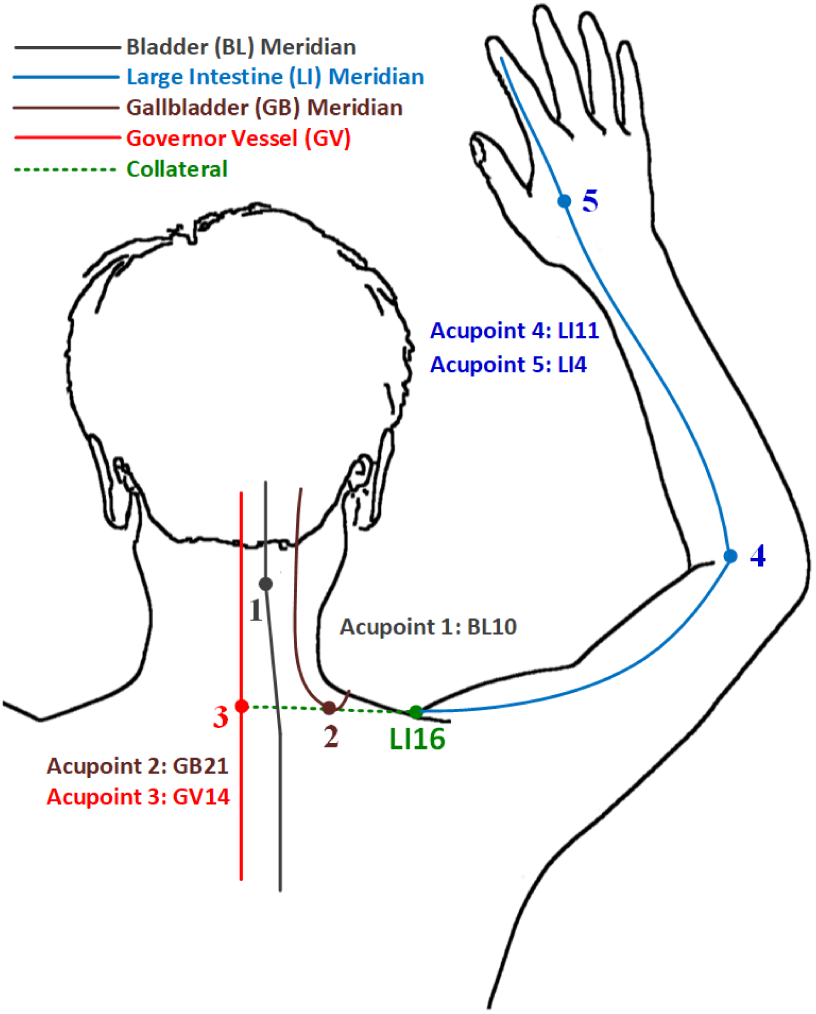
Confluence of longitudinal (Jing) and collateral (Luo) meridians and needle sites: Bladder meridian (BL), Large Intestine meridian (LI), Gallbladder meridian (GB), Governor Vessel meridian (GV) interconnected through a collateral for clinical test with numbers shown: (#2) Driver at Jianjing GB 21; (#3) Driver at Dazhui GV 14 at the 7th cervical vertebra (10); (#1) Sensor at Tianzhu BL10; (#4) Sensor at Quchi LI 11; (#5) Sensor at Hegu LI4 (11)

We set the electro-acupuncture unit at 10 Hz and connected the pulses to #2 and #3 needles. This pair of needles serves as the driver, converting the electrical signal into mechanical vibration. As shown in Figure 2, the needle #2 is in mechanical resonance at 10 Hz with sharp peak amplitude of 0.22 in arbitrary units. Because the needle vibration does not appear to be in a 2D plane, the peak amplitude is an approximate estimate after the FFT. Further down the arm and hand, the needles #4 and #5 are also in resonance at 10 Hz (Figure 3).

**Fig. 2.**
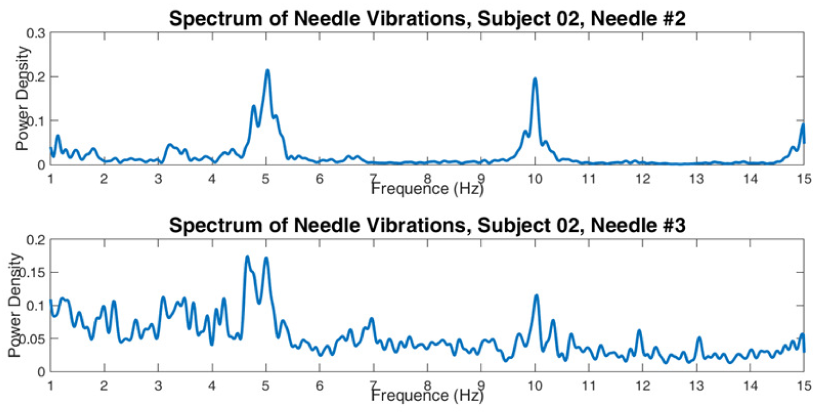
Infrasound frequency spectra of driving signal at acupoints #2 and #3.

**Fig. 3.**
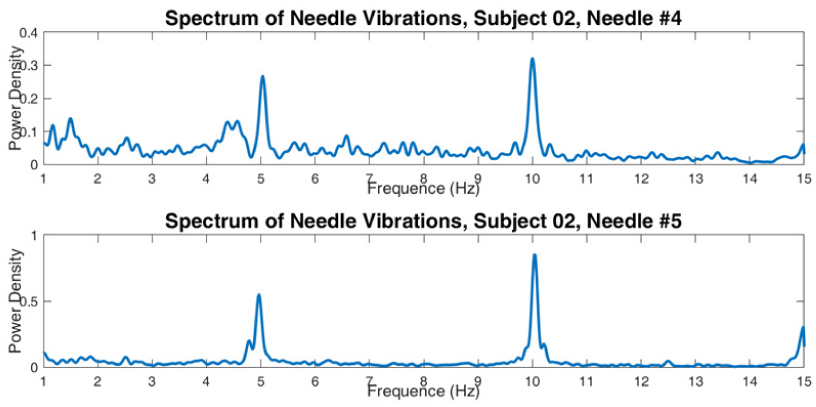
Frequency spectra and resonance of sensing needles at acupoints LI11 (#4) and LI4 (#5). Highest peak is at LI4 (#5) on back of right hand. Note vertical scale variations.

To obtain meaningful statistics, we have taken movies of needle vibration of #2 and #3 as the source drivers and #1, #4, and #5 as three separate receivers with needles inserted into the body as specified by Figure 1. Before electricity is turned on, all 5 needles are scanned and spectra are obtained for each volunteer. These scans constitute the Control group. Afterward, the electricity is turned on, and the same process is repeated to obtain the scans for the Electricity group. Figure 4 shows a typical comparison between the Control (without electricity) and “Electricity” groups. In the Control group data, we have no visible peaks of signal at the source or at the receiving needles. On the receivers with Electricity, #1 needle has only noise but #4 and #5 needles clearly show peak spectra with power gain in needle #5 when compared with the source.

**Fig. 4.**
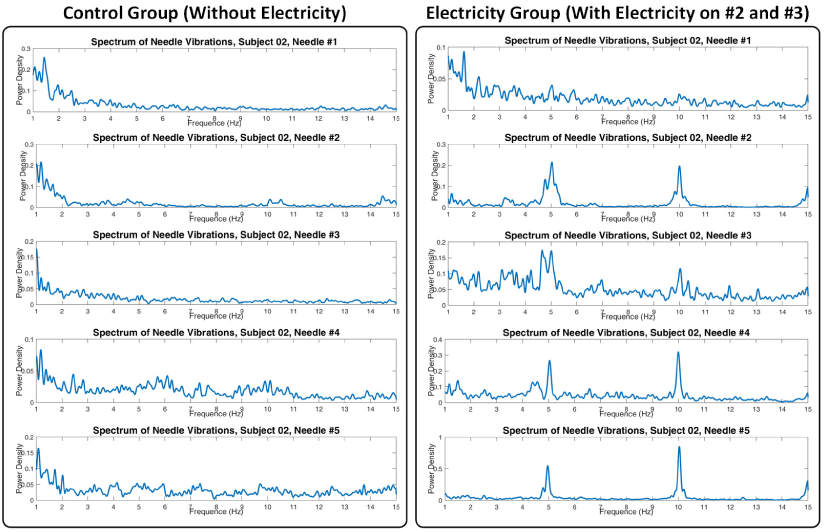
Spectra comparison between Control Group (left column) and Electricity Group(right column). Note vertical scale variations.

Let us next look at what response constitutes positive signaling. Scans of two subjects are off normal as shown in Figure 5. The data on the left column is a broad multi-peak spectrum shown in needle #5. In the right column, there is a positive response with good resonance but a shifted center frequency. We interpret it as an elastic adaptable resonator including a needle implanted in an irregular-shape lump of muscle tissues with anisotropic elasticity. The location of the needle, the depth and angle of penetration, the health and strength of the muscle, and the skill of the acupuncturist all play a role. The mechanical energy from breathing and heart beats and natural adaptation may have modulated or moderated the response as these are unedited raw experimental data.

**Fig. 5.**
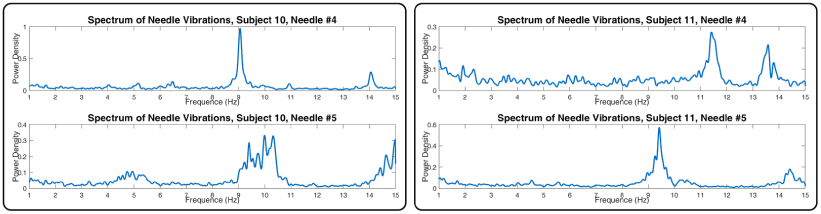
Two examples of spectra comparison of needles #4 and #5: In the left column, the frequency peak of #4 is 9Hz instead of 10Hz and the frequency response is broad-band at around 10Hz for #5. In the right column, the frequency peaks are around 11.5Hz and 9.5Hz for #4 and #5 respectively. Note vertical scale variations.

In the statistical calculation, we postulate that a vibrating frequency observed at the source needles (shoulder and back) is detected at the receiver needles at 50 cm (#4 needle at right elbow) or 80 cm (#5 needle on the back of the right hand) away. Signal transmission is marked as positive, otherwise marked as negative. The Control Group without electrical signal has 10 independent volunteers: one tested as false positive and 9 tested negative. For the Electricity Group, there were 12 independent volunteers: 7 tested positive and 5 tested negative. We have formed the 2-by-2 contingency table (Table 1):

**Table 1.** Summary of responses at needle #4 or #5

|  | Positive | Negative |
| --- | --- | --- |
| Electricity Group | 7 | 5 |
| Control Group | 1 | 9 |

NULL HYPOTHESIS: The positive rate in the “electricity group” is the same as the “control group”, which means there is no non-random association between frequencies detected and electrical acupuncture. ALTERNATIVE HYPOTHESIS: The positive rate in the “electricity group” is higher than that of the “control group”. By using one-tail “Fisher’s Exact Test”, we calculate the p-value from the contingency table: p-value = 0.025. Null Hypothesis is rejected.

Similar to the previous section, we observed the receiver needle #1 that is 10 cm away, relatively nearby. The Control Group has 10 subjects where none of them has positive response. For the Electricity Group, there are 12 subjects where 1 of them has positive response. Thus we can again form the 2-by-2 contingency table (Table 2):

**Table 2.** Summary of responses at needle #1

|  | Positive | Negative |
| --- | --- | --- |
| Electricity Group | 1 | 11 |
| Control Group | 0 | 10 |

The Null Hypothesis and Alternative Hypothesis are the same as the previous section. By using one-tail “Fisher’s Exact Test”, we calculate the p-value as 0.5455. Since the p-value is not less than 0.05, the Null Hypothesis cannot be rejected. It means that the positive rate in two groups has no significant difference. In Figure 1, applied electricity on GB21 (#2) and GV14 (#3) activates the collateral and LI meridian, but BL10 (#1) has no response, thus no signal transmission from #2 to #1 needle in spite of its close distance.

In the last 20 years, various methods were used to test the nature of acupuncture that involves connective tissues, activation of calcium ions, adenosine, ATP and mast cells (12— 16). All of these seemed to have come to the same conclusion that the needle performs a mechanical function but no explicit signaling quantified. This letter shows Qi is an acoustic wave in infrasound frequency, an identifiable physical signaling without ambiguity. It also demonstrates a meridian system that allows one-way traffic, unlike bidirectional air ducts or gas pipes. The unidirectional property matches the clinical descriptions given in Huang Di Nei Jing.

The experiment is very simple. The difficulty is in signal processing and noise suppression. In the low frequency range of infra-sound, the body and its environment is extremely noisy We have patient bed, building and ground vibrations in addition to breathing, heart-beat, blood flow, muscular and body movements. People walking, talking or moving all make inaudible noise. The report of Method tells all the details. We found Kalman filtering was necessary. With Kalman’s invention (17–21), we finally achieved high SNR and quality quantitative data. Acupuncture has no science and is not considered scientific relevant. But this letter delivers both science and technology. It appears to be the only quantitative experiment demonstrating acupuncture follows all the scientific laws and statistics.

The first and foremost result of our study is that acupuncture appears to be purely science based and that there is no magical or superstitious element in its mechanism. This is in contrast with some of the traditional Chinese belief that acupuncture is a unique invention beyond science. It also contradicts the western skepticism as presented in Nature (3) or Wikipedia that “Qi is a non-scientific, unverifiable concept” (January 2017). Infra-sound acoustic signaling is not within our normal daily experience since it is not audible and hardly noticeable. Nevertheless, Qi is observable with a video camera although a conversion into its frequency spectrum is necessary to make the evidence accessible on sight. Second, the details of the meridian network within the limit of the study conform to the sketches given by the standard texts in Chinese medicine (10, 11) and that of Huang Di Nei Jing Ling Shu (2). Thirdly, since acupuncture is pure science, its future employment could be quantified, particularly related to its clinical effectiveness. Improved method and fine tuning of the treatment appear now to be possible.

## Method

Given a sequence of acupuncture images, the region of interest (ROI) is first manually extracted by drawing a box that specifies the area of acupuncture needles. Radon transform (RT) is then used to detect line-shaped patterns in the corresponding ROI. The orientation gauged by the RT is further tracked by the Bayesian Kalman filter (BKF) (17, 18) where a dynamic state-space model is formulated with the system state being composed of the angle and angular velocity of the acupuncture needle. The observations in the tracker are obtained by the RT line detector. Lastly, spectrum analysis of the needle vibration is performed by using the Welch’s method (19). Now we describe various components of the proposed needle vibration analyzing system.

### 1. ROI Identification

In order to estimate the needle vibration accurately, ROI is firstly extracted from the entire acupuncture image so that the interference caused by the background linear features can be mitigated. The ROI segmentation is completed by human hands to guarantee the accuracy. More specifically, a bounding box specifying the ROI is required to be drawn by the user from the first frame. The coordinates of the ROI will then be kept unchanged and further used for the segmentation for following frames, which is reasonable as the needle vibrations during the acupuncture treatment, is usually restricted in a narrow area.

### 2. Radon Transform

After ROI extraction, Canny edge detector (20) is firstly implemented to detect edge features in the ROI. RT is then employed for line detection in the edge map to capture the needle. Compared to Hough transform, RT does not require edge detection and is less sensitive to background noise due to averaging by means of integration. After the needle line detection from the acupuncture frames, the angle between the detected line and horizontal line can be obtained, which will serve as a measurement of the state in the BKF tracker.

### 3. BKF Tracker

As mentioned earlier, the dynamic changes of needle angle can be formulated as a state-space model, in which the state is composed of the angle and angular velocity of the needle. The measurement observation is the needle angle gauged by RT. With such modeling, the angle estimation problem can be solved using the celebrated Kalman filter (KF)-based algorithms (17, 18, 21–23). Since the probability density functions of the needle angle during vibration may be non-Gaussian distributed, we chose the BKF tracker due to its good performance and efficiency for non-Gaussian system estimation. In the BKF tracker, the state and noise densities are approximated with finite Gaussian mixtures, in which the mean and covariance for each component are recursively estimated using the KF. To avoid the exponential growth of mixture components, an efficient Gaussian mixture simplification procedure is employed to reduce the number of mixture components, which leads to lower complexity in comparing with conventional re-sampling and clustering techniques.

### 4. Frequency Analysis

Given the needle angles tracked by the BKF tracker, the Welch’s method can be directly used for frequency estimation and analysis. After that, the spectrum of needle vibration can be obtained.

## Conclusion

We discovered that Qi is an infrasound acoustical shear wave packet, a mathematical wave function, including all chemicals and molecules it carries along with their attributes. Meridian is the muscle pathway or waveguide for the unidirectional transmission of Qi. Efficient signaling comes from the mechanical resonance but less efficient non-resonant transmission might have also made a contribution. In these experimental data and interpretation, Qi and Meridian, originated from the Huang Di Nei Jing, appear to conform to 21st Century Science showing the remarkable medicine appeared understood while practiced clinically 2100 years before our time.

## SI Movies

### SI Movie 1

The video from which the needle vibration spectrum of “electricity group - subject 02, needle #4” (Figure 4, right column, second graph from bottom) is derived from

### SI Movie 2

The video from which the needle vibration spectrum of “electricity group - subject 02, needle #5” (Figure 4, right column, bottom graph) is derived from

### SI Movie 3

The video from which the needle vibration spectrum of “control group - subject 02, needle #4” (Figure 4, left column, second graph from bottom) is derived from

### SI Movie 4

The video from which the needle vibration spectrum of “control group - subject 02, needle #5” (Figure 4, left column, bottom graph) is derived from

## Notes

There is no conflict of interest for all authors. The project has no outside funding.

